# MEG frequency tagging reveals a grid-like code during covert attentional movements

**DOI:** 10.1101/2023.01.29.526079

**Authors:** Giuliano Giari, Lorenzo Vignali, Yangwen Xu, Roberto Bottini

## Abstract

Grid cells in the entorhinal cortex map regular environments with a 60° periodicity, encoding body- and eye-movements’ direction during spatial exploration. Recent evidence in monkeys suggests that grid cells firing is elicited by spatial attention even without eye movements. We investigated whether movements of covert attention can elicit grid-like responses in humans by concurrently recording MEG and eye-tracker. To obtain a measure of grid-like signal non-invasively, we developed a new paradigm based on frequency tagging. While keeping central fixation, participants were presented visually with sequences of linearly-spaced trajectories (15° or 30° in different conditions) formed by static lines or moving dots in separate sessions. Trajectories appeared sequentially on the screen at a fixed rate (6 Hz), allowing different spatial periodicities (e.g., 90°, 60°, 45°) to have corresponding temporal periodicities (e.g., 1, 1.5, 2 Hz), thus resulting in distinct spectral responses in the MEG signal.

Analyses of inter-trial coherence evidenced a higher steady-state response for the frequency corresponding to 60° periodicity compared to control periodicities. This effect was localized in medial-temporal sources and not in control regions. Moreover, in a control experiment using a recurrent sequence of letters featuring the same temporal periodicity but lacking spatial structure, the 60° effect did not emerge, suggesting its dependency on spatial movements of attention. We report the first evidence, in humans, that grid-like signals in the medial-temporal lobe can be elicited by covert attentional movements. Moreover, we propose a new neuroimaging paradigm based on frequency tagging to study grid-like activity non-invasively.

## INTRODUCTION

Understanding the surrounding environment is fundamental for animals’ survival. To this end, sensory information gets organized in so-called “cognitive maps”, an internal model of the environment that supports flexible behavior [Tolman, 1948].

Cognitive maps are thought to be instantiated at the neural level through several neurons responding to spatial variables [Moser et al., 2017]. Among these, grid cells in the entorhinal cortex exhibit multiple firing fields that cover the navigable surface with a 60° rotational symmetry [Hafting et al., 2005]. The finding of spatially modulated cells has been pioneered in rodents, and comparable evidence has been reported in humans in virtual navigation tasks, with invasive direct neural recordings [Jacobs et al., 2013; Nadasdy et al., 2017] but also using non-invasive functional magnetic resonance imaging (fMRI) [Doeller et al., 2010].

Interestingly, in primates the neural mechanisms that evolved to support spatial exploration through locomotion seem to be recruited when space is explored through eye movements [Wirth & Rolls, 2018; Nau et al., 2018]. In non-human primates, grid-cells exhibit their hexagonal firing also when space is explored through saccadic eye movements [Kilian et al., 2012; Meister and Buffalo, 2018]. Similarly, Staudigl and colleagues [2018] reported in humans a higher gamma-band power (60 to 120 Hz) in the medial-temporal lobe (MTL) for saccades aligned to the participants’ grid. Moreover, grid-like responses in the human MTL have been reported by other studies using fMRI during visual search [Julian 2018] and smooth pursuit [Nau et al., 2018]. Findings of a grid-like response during visual exploration suggest the possibility of an attentional mechanism taking place [Bicanski & Burgess 2019]. Gaze position can be conceived as being the overt index of the currently attended location. Attention, however, can also be covertly deployed to peripheral spatial locations without moving the eyes. Interestingly, a grid-like response in entorhinal cells has been reported in non-human primates trained to maintain central fixation while paying attention to a dot moving at random in the periphery [Wilming et al., 2018]. However, in humans there is no evidence of grid-like responses being dissociated from eye movements.

We set out to investigate whether grid-like coding can be elicited, in humans, by movements of covert attention using frequency tagging (FT) [Norcia et al., 2015; Tononi et al., 1998]. Instead of relying on currently established non-invasive methods to detect grid-like responses [see Stangl 2019 for a review] we developed a new method that enables to obtain an objective index of neural response that does not require participants’ overt behavior [Norcia 2015]. In fact, FT relies on the brains’ ability to track regularities embedded within a periodic stimuli presentation, offering the unique advantage of measuring periodic neural responses with high signal-to-noise ratio.

Our FT method relied on rhythmic visual presentation of trajectories, appearing in fixed sequences of angles linearly spaced by either 15° or 30° in different conditions (Fig. 1A). This sequential “clock-like” presentation allowed to embed multiple spatial periodicities at different temporal intervals within the sequence. For instance, in the 30° condition, trajectories separated by 60° (grid-cells periodicity) appeared at 3 Hz, while trajectories separated by 90° (control periodicity) appeared at 2 Hz. The aforementioned frequencies were thus “tagged” with different spatial periodicities (Fig. 1B). If any of these regularities were being tracked, a response at the corresponding frequency would emerge in the MEG signal. As compared to current non-invasive methods, FT does not require to estimate grid orientations with the maximal periodic firing, in that recurrent regularities allow entrainment to each participants’ orientation. We recorded magnetoencephalography (MEG) and eye-tracker while participants attended to the FT presentation. We quantified the neural tracking of the spatial periodicities using inter-trial coherence (ITC) [Ding & Simon, 2013], and observed a grid-like response: The frequency tagged with the 60° periodicity shows higher ITC than control periodicities. At sensor level this effect was found in clusters encompassing occipital and temporal sensors whereas at source level this effect was found specifically in the MTL. Concurrently recorded gaze-location and participants’ accurate performance allowed to ascribe the observed effect to covert attention. In a control experiment we used the same FT design with non-spatially structured stimuli. We observed a different response profile, indicating the dependency of the effect observed in the spatial experiment on covert movements of attention.

**Figure 1.**
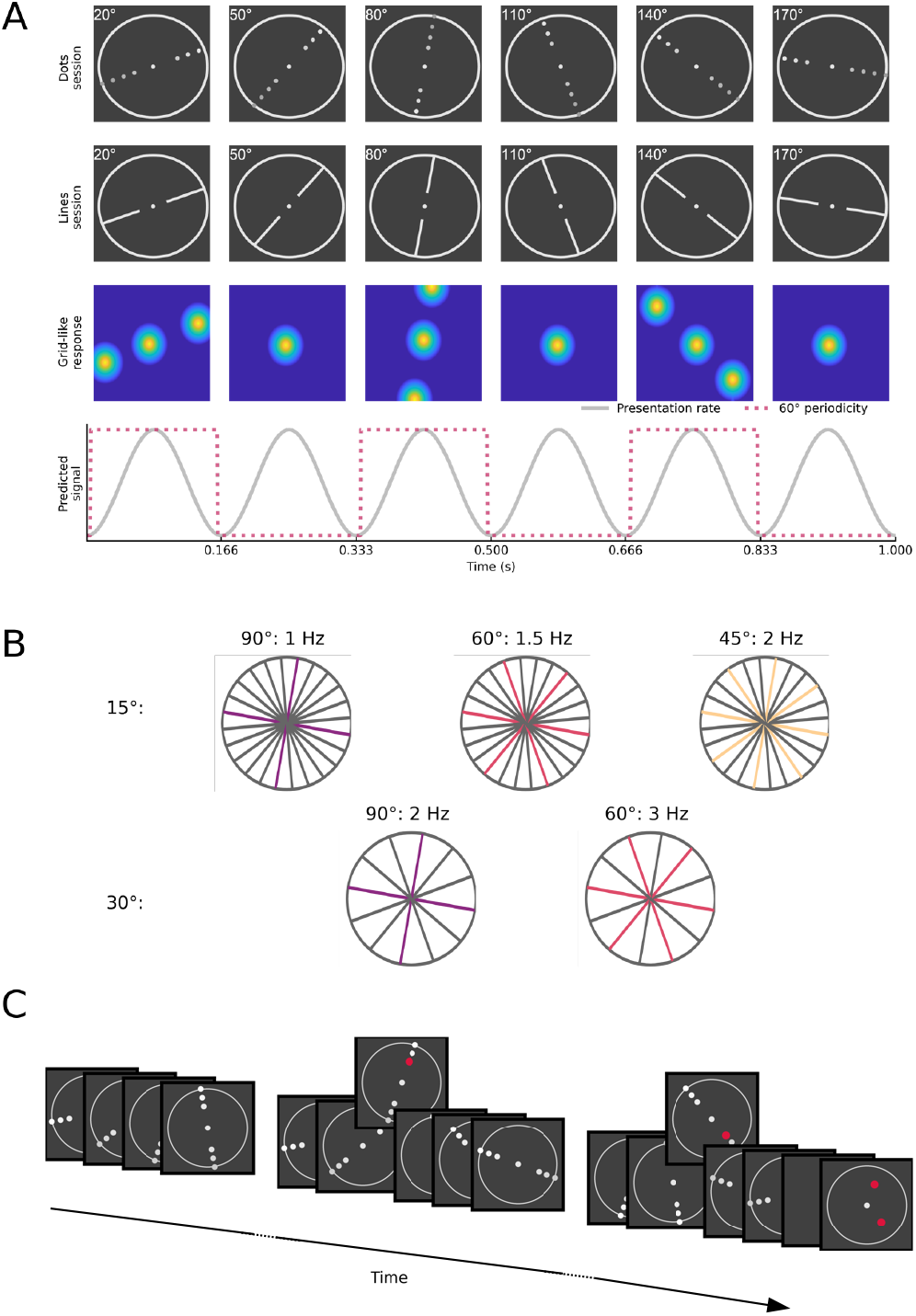
Frequency tagging design. A) Example of a sequence at 30° resolution. Individual trajectories were presented continuously every 0.166 s (6 Hz). The same trajectories were presented in one session as dots moving from one side to the other of the circular arena and in another session as static lines. Crucially, within this sequential presentation were embedded additional spatial regularities. For instance, trajectories separated by 60° appear every 0.333 s (3 Hz). This rhythmic presentation should entrain grid cells that will respond with the same timing. B) Multiple spatial regularities can be embedded within the sequential presentation of linearly-spaced trajectories. Specifically, spatial regularities embedded in the 15° resolution include 90°, appearing every 1 s (1 Hz), 60°, appearing every 0.666 s (1.5 Hz) and 45°, appearing every 0.5 s (2 Hz). The 30° resolution instead include 90°, appearing every 0.5 s (2 Hz) and 60° appearing every 0.333 s (3 Hz). The 60° periodicity corresponds to the grid-like periodicity, whereas the others act as control periodicities. Having two angular resolutions allows to “tag” different frequencies with the same spatial periodicity: trajectories separated by 60° occur at 3 Hz in the 30° resolution and at 1.5 Hz in the 15° resolution, enabling an estimation of the neural tracking that is not tied to a specific chosen frequency. C) Task. During the sequence (both sessions) two red dots appeared for 0.025 s in random locations along the trajectory. To ensure spatial attention, timing of their appearance was randomized but constrained between 45% and 70% for the first dot and between 85% and 90% for the second dot. At the end of the trial, two red dots were presented again and participants had to indicate whether their position is consistent with the one they were keeping in memory.

## RESULTS

### Participants were covertly tracking the spatial trajectories

Twenty-three healthy volunteers completed two MEG recording sessions. In one session, trajectories were formed by dots moving from one side to the other in a circular arena, similar to Nau et al., 2018. In another session, the same trajectories were presented as static lines. Participants were instructed to fixate at the center of a screen while attending to the trajectories. To ensure they were paying attention to the stimuli, they were asked to perform a location memory task (Fig. 1C). This consisted of two red dots appearing at random positions along the trajectories and at random times within the trial sequence. Participants had to remember their exact position, and their memory was tested at the end of each trial. Performance was overall accurate (Fig. 2A), except for one participant that was 2SD below the group mean in both sessions and was thus excluded from further analyses.

**Figure 2.**
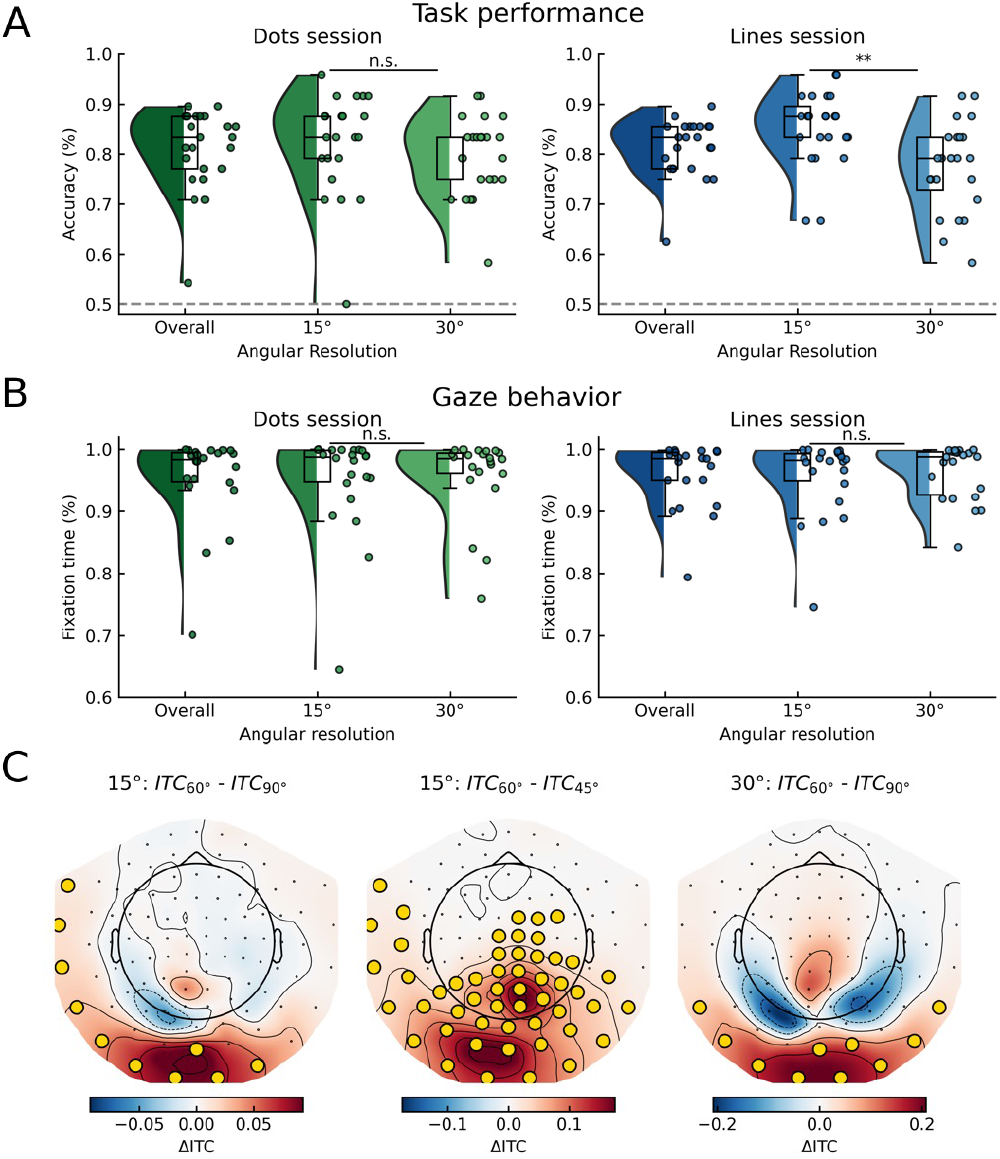
Covert tracking of spatial trajectories elicited a grid-like response detectable with FT. A) Accuracy in the location memory task, averaged over angular resolutions as well as separately for each angular resolution. Performance was overall good (dots session: M=80%, SD=7%, lines session: M=81%, SD=5%). After separating the responses in the two angular resolutions, we observed an higher accuracy in the 15° resolution (t(22)=3.18, p=0.004) in the lines session, while no difference was observed in the dots session (t(22)=1.71, p=0.099). B)Fixation time during the trials, expressed as percentage, averaged over angular resolutions as well as separately for each angular resolution. Participants kept central fixation (4.5° visual angle from the center of the screen) for the majority of the time during the trial. No differences in fixation behavior were found between angular resolutions (dots session: t(21)=−1.06, p=0.300, lines session: t(21)=−1.16, p=0.257). Green shades indicate dots session, Blue shades indicate lines session. Lines above data points indicate significance (n.s.: not significant; *: p < 0.05; **: p < 0.01; ***: p < 0.001) C) Significant clusters at sensor-level in which ITC_60°_ is greater than control ITCs, indicating a grid-like response in occipito-temporal sensors. In the 15° resolution ITC_60°_ is greater than ITC_90°_ (left) and ITC_60°_ is greater than ITC_45°_ (center). In the 30° resolution ITC_60°_ is greater than ITC_90°_ (right). No clusters were found in which control periodicities were higher than 60° periodicity.

Gaze-location data confirmed that participants were keeping central fixation (4.5° visual angle, Wilming et al., 2018) throughout the trial (Fig. 2B). To further make sure that any MEG response can be ascribed to covert attention, trials in which fixation was maintained for less than 80% of the time were excluded from further analyses.

Taken together, participants’ behavior as reflected in task performance and gaze position indicates successful covert tracking of the spatial trajectories.

### MEG FT detects a grid-like response in humans during covert attentional movements

As a first step, we sought to understand whether the FT method was able to detect a grid-like response by quantifying the neural tracking of the spatial periodicities with ITC at sensor-level, an unbiased estimate of the signal as recorded by MEG but with limited spatial precision. After standard MEG preprocessing, the time series of each trial was divided into shorter segments, which then underwent a semi-automatic artifact rejection procedure (see methods). Individual segments were decomposed in the frequency domain with a Fast-Fourier Transformation (FFT), and this complex representation was used to calculate ITC following previous studies [Henin 2019; Ding 2016; Ding & Simon, 2013]. Having observed no statistical differences in the ITC response between the dot and lines sessions in the predicted source level analysis, we averaged the results and carried out our main analyses on this averaged ITC value (see later for the formal comparison and supplementary materials for the individual sessions).

The ITC of the frequencies corresponding to the 60° periodicity (ITC_60°_) was compared to the ITC at the frequency of the control periodicities (ITC_90°_ and ITC_45°_) with a two-sided cluster-permutation test [Maris & Oostenveld 2007], separately for each angular resolutions. We focused on magnetometers given their higher sensitivity to deep sources as compared to gradiometers [Hari 2012]. This analysis revealed significant clusters encompassing occipital and temporal sensors in which ITC_60°_ was higher than the control periodicities (Fig. 2C). Specifically, in the 15° condition, we observed a cluster in which ITC_60°_ was greater than ITC_90°_ (p=0.038 cluster corrected) and a cluster in which ITC_60°_ was greater than ITC_45°_ (p<0.001 cluster corrected). In the 30° condition, we observed a cluster in which ITC_60°_ was greater than ITC_90°_ (p=0.047 cluster corrected). No cluster was found in which control periodicities’ ITCs were greater than ITC_60°_.

This analysis indicates that covert attentional movements in humans also elicit a grid-like response, similarly to non-human primates [Wilming et al., 2018]. The limited spatial resolution of sensor-level analyses localized this response in occipito-temporal sensors, with similar topographies giving rise to the grid-like effect in both angular resolutions (Fig S5B).

### Grid-like response was localized in medial-temporal sources

We performed source localization to investigate which brain areas were responsible for the grid-like effect observed at sensor-level. We used a linearly constrained minimum variance beamformer [Van Veen et al., 1997] to reconstruct the time series of brain activity in each voxel and computed the ITC at the frequencies corresponding to the tagged spatial periodicities (see methods).

We focused our analysis on the MTL (Fig 3A, left), given the a priori hypothesis of its involvement in the generation of the 60° periodic response. The average ITC value in this region of interest (ROI) was compared across sessions with a two-way repeated measure analysis of variance (ANOVA) with factors session (dots, lines) and periodicity (15° resolution: 90°, 60°, 45°; 30° resolution: 90°, 60°) to evaluate whether the neural tracking of the spatial periodicities is elicited differently by moving dots or static lines. This analysis was repeated for each hemisphere and each angular resolution. We found no significant periodicity x session interaction (15°: left: F(2,42)=2.42, p=0.101; right: F(2,42)=1.44, p=0.248; 30°: left: F(1,21)=1.40, p=0.25; right: F(1,21)=0.19, p=0.66). The ITC values of the two sessions (dots and lines) were thus averaged together and used for further analyses (see supplementary materials for the individual sessions results).

**Figure 3.**
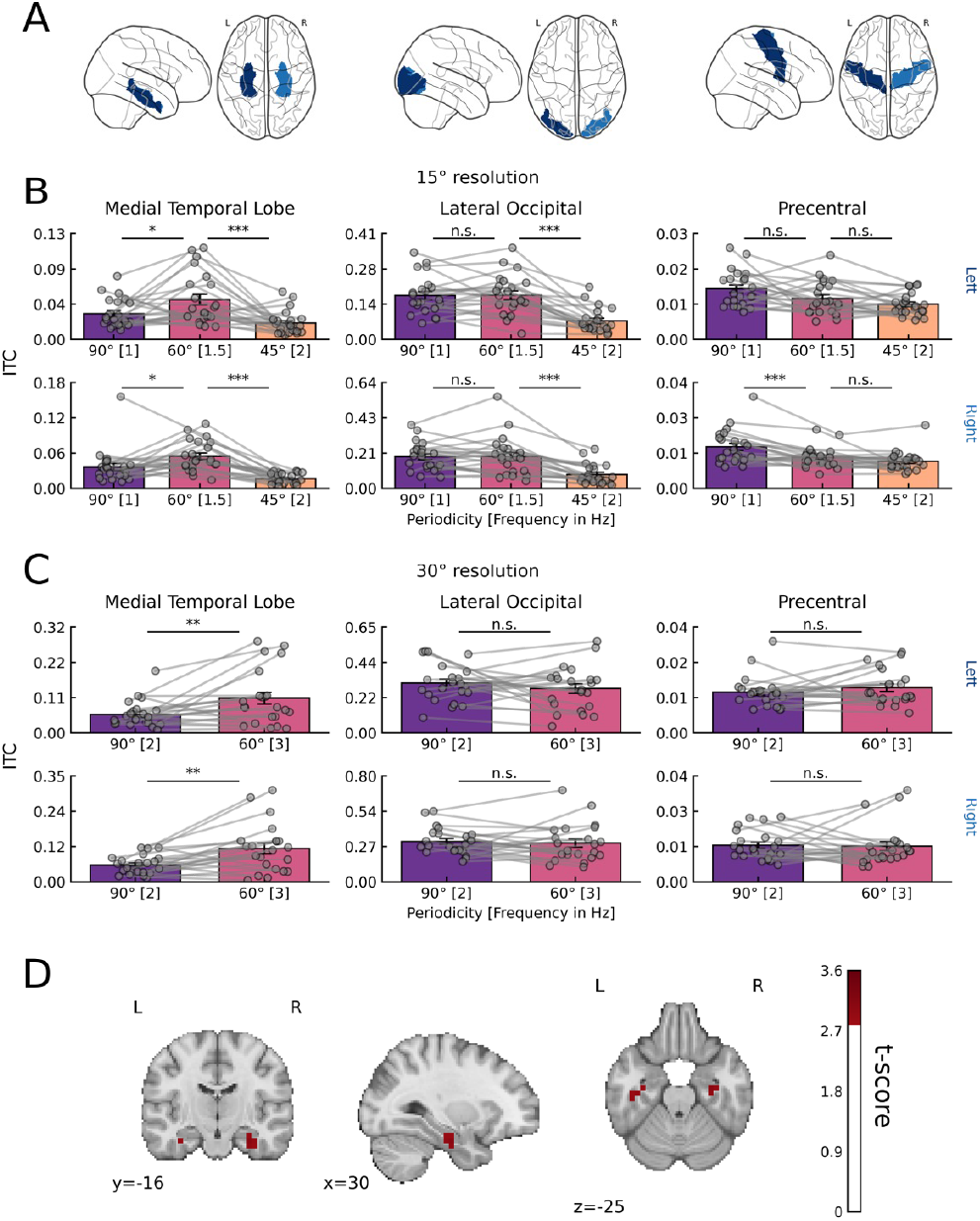
Grid-like response during covert attention originated in the MTL. A)Regions of interest selected for the source level analyses. From left to right: medial-temporal lobe (hippocampus, entorhinal and parahippocampal cortices), lateral occipital and precentral. Dark blue indicate left hemisphere, light blue indicate right hemisphere. B)ITC in the ROIs for each frequency tagged with spatial periodicities at 15° resolution demonstrate the presence of a grid-like response, i.e., the frequency tagged with the 60° spatial periodicity (pink) was higher than both control periodicities (90°: purple, 45° yellow) in the MTL (left) in both the left (top) and right hemisphere (bottom). The grid-like effect was specific to the MTL in that in neither the lateral occipital (center) nor precentral (right) control ROI the 60° periodicity was higher than both control periodicities. C)ITC in the ROIs for each for each frequency tagged with spatial periodicities at 30° resolution demonstrate the presence of a grid-like response in the MTL ROI (left) in both the left (top) and right (bottom) hemisphere, with the 60° periodicity (pink) being higher than the 90° periodicity (purple). This effect was specific to the MTL in that neither the lateral occipital nor precentral ROI show a comparable periodicity preference. Gray dots indicate individual subjects. Error bars indicate standard error of the mean. Lines above data points indicate significance (n.s.: not significant; *: p < 0.05; **: p < 0.01; ***: p < 0.001). D)Conjunction analysis (15°: ITC_60°_ > ITC_90°_ & ITC_60°_ > ITC_45°_ & 30° ITC_60°_ > ITC_90°_) at the cortical level (p<0.005 uncorrected) demonstrated the specificity to the MTL of the grid-like effect.

Most interestingly, the ANOVA identified a main effect of periodicity in both hemispheres, and in both the 15° resolution (left: F(2,42)=9.27, p<0.001; right F(2,42)=16.46, p<0.001) and 30° (left: F(1,21)=8.63, p=0.008; right F(1,21)=11.89, p=0.002). Planned paired t-test indicated that in the 15° resolution, ITC_60°_ was significantly greater than both ITC_90°_ (left: t(21)=2.35, p=0.028; right: t(21)=2.45, p=0.022) and ITC_45°_ (left: t(21)=3.92, p<0.001; right: t(21)=6.80 p<0.001) and in the 30° resolution ITC_60°_ was significantly greater than ITC_90°_ in both the left (t(21)=2.92, p=0.008) and the right MTL (t(21)=3.43, p=0.002). These results suggest the presence of a grid-like response in the MTL during covert attentional movements.

To investigate whether the grid-like effect was specific to the MTL, we compared the MTL with two control ROIs, the lateral occipital and precentral cortices (Fig 3A, center and right). We ran a two-way repeated measures ANOVA with factors ROI (MTL, lateral occipital, precentral) and periodicity (15° resolution: 90°, 60°, 45°; 30° resolution: 90°, 60°), separately for each hemisphere and each angular resolution. We found a significant two-way interaction in the 15° condition (Fig 3B) in both the left (F(2.51, 52.75)=29.62, p<0.001, Fig. 3A) and right hemisphere (F(2.17, 45.66)=20.82, p<0.001, Fig. 3B). Post-hoc t-test indicated that the effect was indeed specific to the MTL as in neither the lateral occipital nor the precentral ROI, ITC_60°_ was greater than both control periodicities (lateral occipital: left: ITC_60°_ vs ITC_90°_: t(21)=0.07, p=0.937; ITC_60°_ vs ITC_45°_: t(21)=6.43, p<0.001; right: ITC_60°_ vs ITC_90°_: t(21)=0.06, p=0.949; ITC_60°_ vs ITC_45°_: t(21)=6.09, p<.001; precentral: left: ITC_60°_ vs ITC_90°_: t(21)=−1.93, p=0.066; ITC_60°_ vs ITC_45°_: t(21)=1.83, p=0.080; right: ITC_60°_ vs ITC_90°_: t(21)=−4.15, p<0.001; ITC_60°_ vs ITC_45°_: t(21)=1.82, p=0.081). In the 30° resolution (Fig. 3C) we observed a significant two-way interaction in the left (F(1.35, 28.27)=4.94, p=0.025) and in the right hemisphere (F(1.39, 29.13)=4.79, p=0.026, Fig. 3D). Post-hoc t-tests confirmed that in the control regions there was no significant difference between the ITCs (left: lateral occipital: t(21)=−1.01, p=0.322; precentral: t(21)=1.07, p=0.294; right: lateral occipital: t(21)=−0.39, p=0.698; precentral: t(21)=−0.14, p=0.884).

To localize this effect at the cortical level, we performed a conjunction analysis [Nichols et al., 2005]. This analysis revealed clusters of voxels in the bilateral MTL that survived an uncorrected threshold (p < 0.005) (Fig. 3D).

These results confirm that the grid-like response elicited by covert attentional movements originates in the MTL and was not present in control regions.

### Temporal structure of the FT design cannot explain the grid-like response

Temporal regularities in stimuli presentation constitute the most important feature of FT designs [De Rosa et al., 2022]. When multiple regularities are present (as in our case), their interaction can result in additional neural responses at frequencies corresponding to any sum of the originally tagged frequencies and their multiples (i.e., intermodulation [Norcia et al., 2015; Gordon et al., 2019]). Intermodulation of salient rhythms inherent to the previously presented FT design (e.g., presentation rate and “turn” of the clock-like presentation [Cracco et al., 2021]) could potentially provide an alternative interpretation of the effects that were observed (Fig. S3).

We thus ran a control experiment to rule out the possibility that the grid-like response reported in the previous experiment was the byproduct of other temporal regularities.

Specifically, we designed another FT task having the same temporal regularities (i.e., the repeating sequences) but using non-spatially-structured stimuli (i.e., letters). In the spatial experiment sequences were of different duration in the two angular resolutions, due to a difference in the number of trajectories. We replicated this feature using a different number of letters in the sequences (15°: twelve letters from A to N; 30°: six letters from A to F, see methods and Fig S4).

Data from twenty-two healthy participants that took part in this non-spatial experiment were analyzed following the same approach of the spatial experiment (see methods). If the temporal structure of stimulus presentation and the interaction between multiple frequencies was the cause of the results in the spatial experiment, we should see a preference for the frequency that was previously tagged with 60° periodicity (i.e., 1.5 Hz in 15° resolution and 3 Hz in 30° resolution) in this non-spatial experiment as well. To test this hypothesis, we extracted the ITC of the frequencies that in the spatial experiment were tagged with spatial regularities (Fig. 1B) and compared them at both sensor- and source-level, both within and between experiments.

At sensor-level, a two-sided cluster permutation test (Fig. 4A) in the condition that corresponded to the 15° angular resolution identified an occipito-temporal cluster in which ITC at the frequency corresponding to 60° spatial periodicity (ITC_Cont60°)_ was greater than ITC at the frequency corresponding to 90° (ITC_Cont90°_) (p<0.001 cluster corrected) and two clusters in which ITC at the frequency corresponding to 45° (ITC_Cont45°_) was higher than ITC_Cont60°_, one located in left temporal sensors (p=0.010) and one in right frontal sensors (p=0.005). In the condition corresponding to 30°, we identified an occipito-temporal cluster in which ITC_Cont60°_ was greater than ITC_Cont90°_ (p=0.013 cluster corrected). The condition corresponding to the 15° resolution showed clearly different results as compared to the spatial experiment, with sensors in which ITC_Cont45°_ was greater than ITC_Cont60°_. In the condition corresponding to the 30° resolution, with only one control periodicity, it was more difficult to characterize a grid-like response as compared to a more general response elicited by the temporal structure. Nevertheless, the topographies of the two experiments were different (Fig S5A) indicating different neural origin of the effect. Moreover, we found a high correlation of the grid-like response (i.e., the difference between ITC_60°_ and ITC_90°_) between angular resolutions in the spatial experiment, and not when correlating the corresponding frequencies of the non-spatial experiment (Fig. S5B).

**Figure 4.**
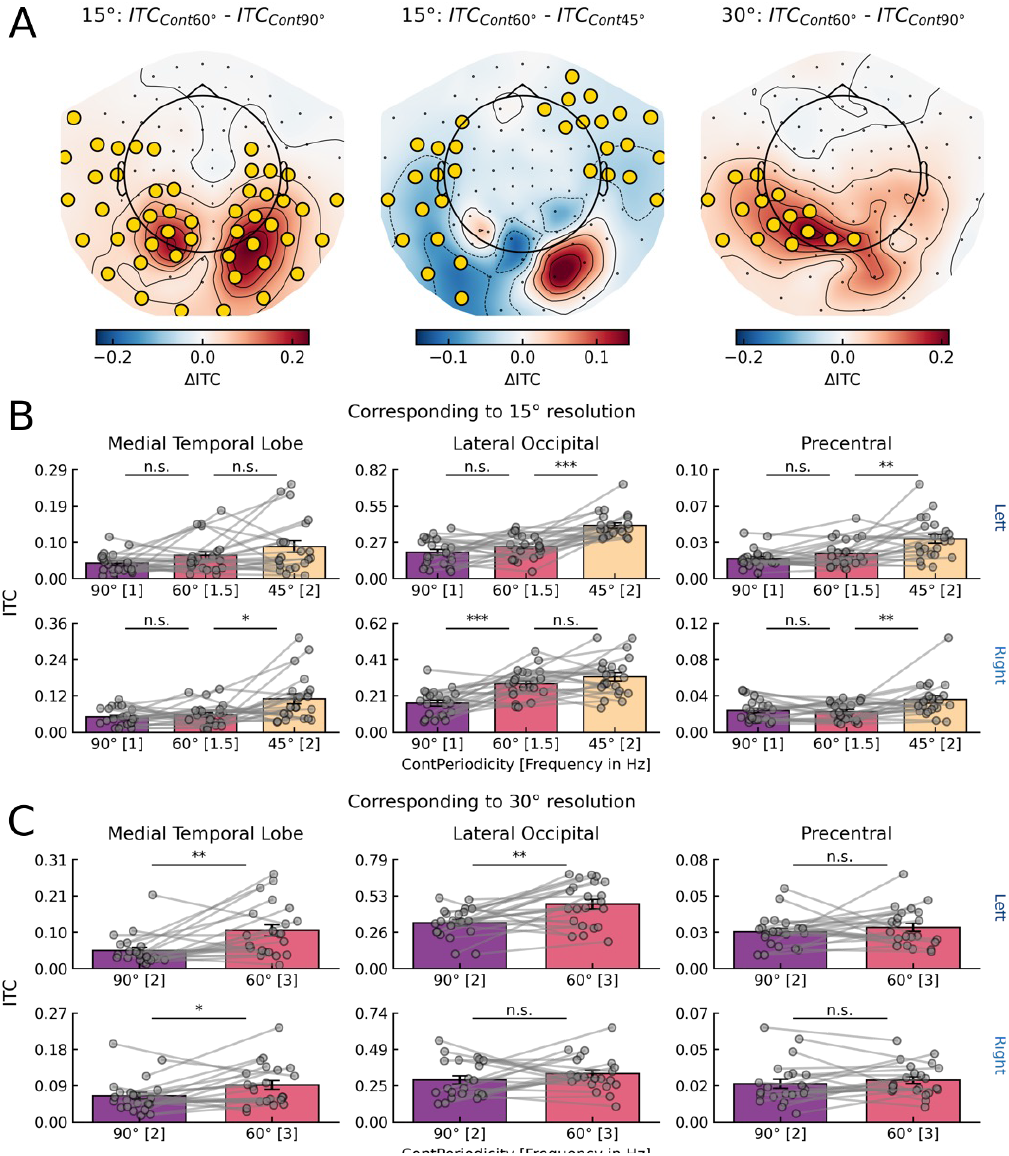
Temporal structure does not elicit a grid-like response. A) Sensor-level clusters observed in the non-spatial experiment comparing the frequencies tagged with spatial periodicities in the spatial experiment. In the condition corresponding to the 15°, ITC_Cont60°_ was greater than ITC_Cont90°_ whereas ITC_Cont60°_ was lower than ITC_Cont45°_. In the condition corresponding to 30° instead ITC_Cont60°_ was greater than ITC_Cont90°_. B) In the condition corresponding to 15° resolution, a significant three-way interaction (experiment x ROI x periodicity) indicates different response profiles between the spatial (Fig 3B) and non-spatial experiment. In the non-spatial experiment, no significant differences between frequencies were identified in the left MTL ROI (top left), while in the right (bottom) MTL ITC_Cont60°_ was significantly lower than ITC_Cont45°_. C) In the condition corresponding to the 30° resolution, a significant three-way interaction (experiment x ROI x periodicity) indicates different response profiles between the spatial (Fig 3C) and non-spatial experiment only in the left (top) and not in the right (bottom) hemisphere. This significant interaction was caused by ITC_Cont60°_ being greater than ITC_Cont90°_ in the left lateral occipital, in that no differences between experiments were found in the MTL. Gray dots indicate individual subjects. Error bars indicate standard error of the mean. Lines above data points indicate significance (n.s.: not significant; *: p < 0.05; **: p < 0.01; ***: p < 0.001).

We then reconstructed the sources of the signal to further explore the effects elicited by the temporal structure and make sure that a 60° preference was not present in the MTL. Finally, we compared the ITC values in the ROIs between experiments using ANOVAs and bayesian model comparison.

In the 15° condition (Fig 4B), a mixed ANOVA with experiment as between-subjects factor and ROI and periodicity as within-subjects factor identified a significant three-way interaction in both the left (F(2.73,114.58)=39.43, p<0.001) and right hemisphere (F(2.77, 116.27)=22.93, p<0.001), indicating different patterns of ROI x periodicity interaction across the two experiments (spatial vs non-spatial). Indeed, in the MTL the ITC_Cont60°_ was never greater than both the control periodicities. Moreover, contrary to the spatial version of the experiment, the frequency preference showed similar trends in both the MTL and the control regions (See Fig 4B and Table S1 for post-hoc pairwise comparison within each ROI). In order to directly compare ITC_60°_ with both control ITCs at the same time, we fitted (within each experiment) linear (L) and quadratic (Q) models to the group-level data and compared their goodness of fit with bayes factor (BF, [Wagenmakers, 2007]). In the MTL ROI (Fig 5A), we found very strong evidence [Raftery, 1995] for the quadratic model centered on the 60° periodicity over the linear model in both the left (BF_QL_=150.05) and right hemisphere (BF_QL_=2203.92) in the spatial experiment. Conversely, in the non-spatial experiment, we found positive evidence for the linear model over the quadratic model for both the left (BF_LQ_=8.07) and weak evidence in the right hemisphere (BF_LQ_=2.04). This is confirmed at the cortical level, where the right MTL shows very strong evidence in favor of the quadratic model as compared to the linear model in the spatial experiment (Fig. 5B).

**Figure 5.**
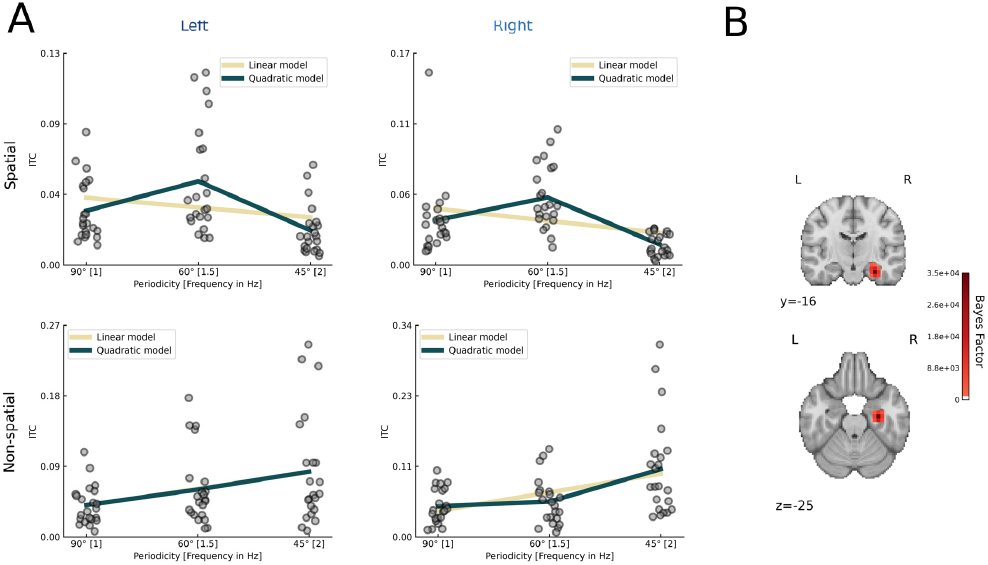
Bayesian model comparison identifies a quadratic trend in the spatial experiment and a linear trend in the non-spatial experiment. A) In the spatial experiment (top), there was very strong evidence in favor of a quadratic model centered on ITC_60°_ as compared to a linar model in both the left (BF_QL_=150.05) and right hemispehre (BF_QL_=2203.92). Conversely, in the non-spatial experiment, there was positive evidence for a linear model as compared to a quadratic model in the left hemisphere (BF_LQ_=8.07) while weak evidence for the linear model in the right hemisphere (BF_LQ_=2.04). B)Voxel-wise model comparison in the spatial experiment identified very strong evidence for the quadratic model as compared to the linear model in the MTL.

In the 30° resolution (Fig 4C), we found a significant three-way interaction (experiment x ROI x periodicity) in the left (F(1.3, 54.7)=11.14, p<0.001) but not in the right hemisphere (F(1.23, 51.84)=2.56, p=0.109). Post-hoc comparisons (see Fig. 4C and Table S1) revealed that a preference for the frequency that was previously tagged with 60° spatial periodicity was found in the bilateral MTL but, contrary to the spatial experiment, also in a control region, the left lateral occipital.

Taken together, these results demonstrated that the FT temporal structure generated different neural responses based on the task. The presentation of spatial sequences and their covert attentional tracking resulted in a grid-like response specific to the MTL. The same temporal structure but no spatial allocation of attention instead produced different response profiles across the whole brain. In the 15° condition, we observed a linear increase of ITC with the frequency. In the 30° condition it was more difficult to characterize the grid-like response as independent from other temporal regularities, given the presence of a single control periodicity. Nevertheless, differences in both topographies and control regions make the MTL grid-like effect specific to the spatial experiment, suggesting differences in the mechanisms that gave rise to the observed response in the non-spatial experiment. In this respect, the left lateralization may be suggestive that lexical regularities (that have been shown to be detectable with FT [Lochy 2015; Lochy 2018]) were playing a role.

## DISCUSSION

We demonstrated that a grid-like response, i.e., a preference for spatial trajectories aligned with a 60° periodicity as compared to control periodicities, was elicited in the human MTL by movements of attention. This effect arises independently from overt oculo-motor behavior, providing the first direct evidence that also in humans covert attention induces grid-like responses.

To do this, we used MEG and eye-tracker in combination with a newly developed method based on frequency tagging. The proposed method relies on the periodic visual presentation of spatial trajectories. Crucially, unbeknownst to participants, these trajectories exhibited spatial regularities corresponding to the periodic firing of grid cells. This resulted in a grid-like response that was specific to the medial temporal lobe, thus demonstrating the biological plausibility of the cellular-level signal that was picked up non-invasively with MEG. Moreover, in a control experiment, we provided evidence that the grid-like signal depended on covert movements of attention, and not on the temporal structure of our task.

### Covert attention elicited a grid-like signal in the human MTL

Previous literature had already demonstrated that overt visual exploration can result in a grid-like response in the MTL [Nau 2018; Killian 2012; Julian 2018; Staudigl 2018] strengthening the relationship, in primates, between the neural mechanisms supporting spatial exploration through bodily and ocular movements [Rolls & Wirth 2018; Nau 2018]. Gaze location is often considered a proxy for spatial attention. But attention can also be covertly moved in space independently of eye-movements, resulting in a similar grid-like response in non-human primates [Wilming et al., 2018]. Here we demonstrated that a grid-like response was present also in humans while attention was covertly deployed to spatial locations.

By replicating findings from non-human primates, we provided comparable evidence across species for an attentional mechanism being able to give rise to grid-like responses. The direct demonstration of the presence of a mechanism for covert exploration in humans can account for the numerous findings of grid-like responses during imagined navigation [Horner 2016; Bellmund 2016] as well as navigation of conceptual spaces [Bellmund et al., 2018; Bottini & Doeller, 2020]. Moreover, the identified mechanism is in line with findings of memory-guided attention, suggesting that the MTL participates in guiding attention towards behaviorally relevant spatial locations to support future perception and action [Gunsely & Aly 2020; Stokes et al., 2012; Summerfield 2006]. The outcome of this MTL-based attentional orienting could then become visible at both the neural and behavioral level through medial-temporal interactions with the oculomotor system [Buffalo & Meister, 2016; Ryan & Shen 2019], leading to memory-guided oculomotor behavior [Kragel & Voss 2022].

### Frequency tagging as a non-invasive tool to assess the grid-like response

A second purpose of the present study was to develop a novel, non-invasive method that allows detection of the grid-like response. The seminal paper by Doeller and colleagues [2010] opened the possibility to study the proxy of a cellular response non-invasively giving rise to numerous discoveries on the functioning of grid-like firing in humans. While this method has been used successfully in both healthy participants and special populations [e.g., Kunz 2015; Bierbrauer 2020], it relied on participants’ compliance to perform a task and undergo long training procedures. This greatly limited the possibility to investigate grid-like signals with populations that instead may have problems in performing cognitive tasks, such as patients, or may not be able to undergo long experiments, such as children.

We developed a method to elicit a grid-like response based on FT that overcomes these limitations and can be used to further advance our understanding of grid-like responses in the human brain. FT in fact has been employed successfully with special populations [e.g., De Heering & Rossion, 2015; Buiatti et al., 2019; Vettori et al., 2020]. Moreover, FT can be used to study high-level cognitive processes, such as attention [Müller et al., 2003; Joon Kim et al., 2007] or understanding spatial relations [Adibpour et al., 2021].

Taken together, our results indicate that the proposed method is a valid alternative to the now-standard non-invasive analytical approaches to detect a grid-like signal, with the advantage of requiring less effort from the participants and thus being potentially useful with special populations.

## Supporting information

Supplementary results

## EXPERIMENTAL MODEL AND SUBJECT DETAILS

Twenty-four participants (12 male, age M=25,88 years; SD=4,84) were recruited to participate in the spatial experiment that consisted in two magnetoencephalography (MEG) sessions, conducted in different days (maximum eight days apart). One participant did not show up for the second session and was excluded from the analysis. Another sample of twenty-four participants (9 male, age M=23,91 years; SD=3,51) were recruited to participate in the non-spatial experiment.

All had normal or corrected to normal vision and no history of neurological disorders. Prior to each session they gave written informed consent to participate in the experiment. All procedures were approved by the ethical committee of the University of Trento.

## METHOD DETAILS

### Spatial experiment design

The experiment consisted in the visual presentation of spatial stimuli (trajectories). These were defined dividing a circle in 24 and 12 equidistant points, resulting in two angular resolutions (15° and 30°) that were presented in different trials. These points were rotated 10° clockwise to avoid trajectories to appear along the cardinal axes. Each trajectory connected two opposite points (i.e., 180° apart) in the circle.

In separate recording sessions these trajectories were presented to the same participants either as a sequence of dots, moving from one end to the other of the circle, or as static lines, covering the whole trajectory at once. The order of the sessions was counterbalanced across participants. Note that static lines lack directionality, effectively reducing the number of trajectories to half. That is, a trajectory starting at 20° and ending at 200° occupied the same portion of space as a trajectory starting at 200° and ending at 20°. In the dots session half of the trajectories were presented as starting from the opposite side of the circle, effectively making opposite trajectories indistinguishable between each other. The total number of trajectories is thus 12 in the 15° resolution and 6 in the 30° resolution.

Stimuli were generated using MATLAB (version 2012b, The Mathworks, Natick, MA, USA) and presentation was controlled using PsychToolbox [Brainard 1997].

A new trajectory was presented every 0.166 s (6 Hz) using a typical frequency-tagging (FT) approach. In each trial were presented 264 trajectories, for a total duration of 44s. These appeared sequentially, in a clock-like fashion (e.g. 20°, 50°, 80°, 110° etc… in the 30° resolution, Fig. 1A), with clockwise/counter-clockwise direction balanced across participants. Crucially, the periodic presentation of regularly spaced trajectories allowed to embed multiple spatial periodicities in the trajectories’ sequence, each appearing at fixed and distinct temporal intervals such that each spatial periodicity “tags” a unique frequency (Fig 1B). Specifically, at 15° resolution, trajectories separated by 60° appear every 0.666 s (1.5 Hz), trajectories separated by 90° every 1 s (1 Hz) and trajectories separated by 45° every 0.5 s (2 Hz). At 30° instead, trajectories separated by 60° appear every 0.333 s (3 Hz) while trajectories separated by 90° appear every 0.5 s (2 Hz). Both 60° and 90° occur in both spatial resolutions, but tagging different frequencies in each. This enables an estimation of their neural tracking that is not tied to a specific chosen frequency as well as direct comparison between responses in the two spatial resolutions.

Multiple sequences with the same trajectories’ order were presented in a trial to allow for entrainment to the spatial periodicities. Specifically, individual sequences lasted 2s (15°) and 1 s (30°), with each sequence containing one instance of each trajectory. With the clock-like presentation one sequence corresponds to half-turn of the clock. The starting trajectory of the sequence was randomized across trials such that each trajectory was used as a start twice (15°) or four times (30°) over the experiment.

Participants were instructed to fixate in the centre of the screen while paying attention to the trajectories that were shown in the periphery. To avoid following saccades, we defined a fixation window of 4.5° of visual angle [Wilming et al., 2018] in which no visual stimulus was presented except for a fixation dot that remained on the screen centre for the whole trial duration.

To ensure participants were covertly tracking the trajectories, we asked them to perform a location memory task (Fig 1C). Two red target dots appeared at random position along a trajectory for 0.025 s. The timing of their appearance was randomized but constrained between 45% and 70% of the trial for the first dot and between 85% and 90% of the trial for the second dot. This timing was chosen to ensure participants were focused until the end of the FT presentation. Participants were instructed to remember the exact position of these two red target dots for future recall. At the end of each trial two red test dots appeared in three possible configurations, each with equal probability: i) test dots occupy the same spatial positions as target dots; ii) only one test dot does not occupy the same position of the target dots; iii) there was no overlap between test and target dots.. Participants had to respond via button-press with the right index finger to indicate same position and with the right middle finger to indicate different position, irrespective of the number of dots being in a different position. The probability of these occurrences was 50%.

The experiment was divided in six blocks of eight trials each, for a total of 48 trials, 24 in each angular resolution. In each block there were four consecutive trials of the same angular resolution, and the resolution presented as first was randomized and balanced across blocks.

The timing of the presentations was controlled by the computer. A 15 s break was included after every trial, while 30 s were allowed after 4 consecutive trials. After each break, a 3 s countdown informed participants that a new trial was about to start. The trial then only started after participants had fixated for at least 500 ms within a 2° visual angle fixation window centred on the fixation dot. At the end of each trial, a warning message appeared on the screen if the participant’s eyes remained outside the fixation window (4.5°) for more than 20% of the overall time. After each block participants were allowed a longer, self-timed, break of about 2 min.

### Non-spatial experiment design

With this control experiment we wanted to test the hypothesis that temporal regularities inherent to the FT design can generate a periodic neural response that is comparable to the grid-like signal measured in the spatial experiment. Temporal regularities are defined here by the number of stimuli presented in the sequence and the individual sequences’ duration, which varied between the angular resolutions of the spatial experiment. We thus reproduced these features using stimuli without spatial structure. Specifically, we reproduced the temporal structure of the 15° resolution by creating a sequence of 12 letters (A to N). Similarly, the 30° resolution was reproduced by creating a sequence of 6 letters (A to F).

Individual letters were visually presented every 0.166 s (6 Hz) with contrast modulation [Lochy 2019; Lochy, 2015]. Other potentially salient temporal regularities that may be tagged by this FT design include the sequence duration. Sequences corresponding to the 15° resolution lasted 2 s (0.5 Hz) while sequences corresponding to 30° resolution lasted 1 s (1 Hz). Frequencies of interest in the spatial experiment are not tagged with spatial regularities anymore. Any effect observed at these frequencies cannot be ascribed to spatial regularities but should be interpreted as arising from the intermodulation of the presentation rate and other potentially salient rhythms such as the sequence rate.

Participants were instructed to fixate in the centre of the screen while paying attention to the letters’ sequences. Moreover, to keep them engaged, twice during a trial and with the same time constraints as in the spatial experiment the fixation dot turned red for 0.025 s, a change that the participants were instructed to promptly detect (max 3 sec) via right index finger button press.

### MEG and eye-tracker acquisition

MEG data were acquired at the Center for Mind/Brain Sciences of the University of Trento with an Elekta Neuromag 306 MEG system (Elekta, Helsinki, Finland), composed of 102 magnetometers and 204 planar gradiometers, placed in a magnetically shielded room (AK3B, Vakuumschmelze, Hanau, Germany). The head-shape of the participants was digitized (Fastrak Polhemus, Inc., Colchester, VA, USA) prior to acquisition in each session, along with fiducial points (nasion, left and right periauricular) and five head position indicator (HPI) coils, three placed on the forehead and one behind each ear. Both fiducials and HPIs were digitized twice to ensure precision (< 2 mm difference).

Before entering the MEG, participants performed a short practice block (4 trials, 2 of each angular resolution) to familiarize with the FT design and the task. They received written instructions before the practice and feedback on their performance after each trial.

Participants sat upright in the MEG chair with their head as close as possible to the dewar. The eye-tracker (Eyelink 1000 Plus, SR Research Ltd., Ottawa, Canada) was positioned to ensure optimal recording of both eyes. A nine-point calibration procedure was carried out before each block.

Continuous MEG data were recorded at 1000 Hz with hardware bandpass filters in the range 0.1-330 Hz. Along with MEG, we also recorded the time series of a photodiode that tracked the colour-change of a small square on the top-left corner of the screen (not visible to the participants). This was used in the analysis to correct for potential delays in the stimuli presentation. Eye-tracker was recorded separately for each trial at a sampling rate of 1000 Hz. Stimuli were projected on a translucent whiteboard, positioned 1 m in front of the participant, using a ProPixx projector (VPixx Technologies, Canada) at a 120 Hz refresh rate. Responses were collected using a MEG compatible button response box (VPixx Technologies, Canada).

After each session a 5 minutes empty-room measurement was recorded to be used for noise modelling in the source reconstruction procedure.

Four blocks from different participants had technical issues and were not included in the final analysis.

### MEG and eye-tracker preprocessing

Raw task data and empty-room MEG time series were visually inspected and artefactual channels (M=7.59, SD=6.75) were marked for exclusion and interpolation through MaxFilter (temporal signal suppression, Taulu & Simola 2006). Raw task data was subsequently realigned to the recording block that minimized the Euclidean distance across blocks, separately for each session.

After application of Maxfilter, continuous raw data were filtered (High pass: 0.1 Hz, Low-pass 40 Hz) and segmented into 44 s–long trials, starting from the onset of the first trajectory. Trial onset was corrected for potential delays using a photodiode.

Hence, for each trial we quantified fixation behaviour by computing the percentage of time the right eye position was within the 4.5° fixation window. Trials in which this metric was below 80% of the overall time were excluded from further analysis (percentage of rejected trials: spatial experiment: dots session: 5.7%; lines session: 5.1%; non-spatial experiment: 7.69%). In the non-spatial experiment one participant was excluded at this stage due to the low number of trials left after exclusion (33%). To ensure consistent timing across trials in the appearance of the spatial periodicities, trials were realigned to the first presentation of the 350° trajectory, reducing their length to 40 s and 42 s in the 15° and 30° resolution, respectively.

To increase the signal-to-noise ratio, as suggested by Benjamin et al., [2021], trials’ time series were further divided in shorter, non-overlapping segments of 8 s and 6 s, resulting in 5 and 7 segments per trial in the 15° and 30° resolution, respectively. Segment duration was chosen for multiple reasons: i) It allows an integer number of repetitions of each individual sequence (4 and 6); ii) each segment contains an integer number, as well as an high number, of cycles of the frequencies of interest and enables as a high frequency resolution (0.125 Hz and 0.167 Hz). Segmented data underwent a semi-automatic artifact rejection procedure in which variance and kurtosis metrics were computed, akin to the visual artifact rejection procedure implemented in the Fieldtrip package [Oostenveld 2011]. Segments above two standard deviations in each individual metric were visually inspected for the presence of artifacts (e.g., remaining channel jumps). This procedure led to the exclusion of an additional 0.68% of segments in the dots session of the spatial experiment and 1.04% of segments in the lines session, while 0.83% of segments were excluded in the non-spatial experiment.

### Frequency analysis

Artifact-free segments were subjected to a fast-fourier transformation (FFT) separately for each channel (at sensor-level) or voxel (at source level). From the complex representation of each segments’ time series in the frequency domain we computed the inter-trial coherence (ITC) as follows:

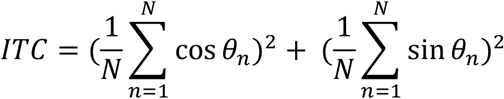

where *θ* is the phase of the individual segment (n) as obtained from the FFT [Henin 2021; Ding 2016; Ding 2013].

This metric quantifies the synchronization of the neural response across segments and ranges from 0 to 1, where 1 indicates perfect synchronization. We then isolated the ITC at the frequencies “tagged” with the spatial periodicities (see “Spatial experiment design”), given the narrowband response afforded by the FT method. For comparison, the same frequencies were selected in the non-spatial experiment. However, these frequencies bore no spatial meaning, as in this control experiment they were not tagged with spatial regularities.

### Source reconstruction

Structural T1-weighted images were acquired at the Center for Mind/Brain Sciences of the University of Trento in a 3T Siemens Prisma scanner (Siemens, Erlangen, Germany) with a Multi-Echo MPRAGE (MEMPRAGE) sequence with the following parameters: FoV = 256mm; Voxel Size = 1×1×1 mm3; TR = 2530 ms; TE1 = 1.69ms; TE2 = 3.55ms; TE3 = 5.41ms; TE4 = 7.27ms and a flip angle of 7°. Two participants of the spatial experiment did not perform the MRI session.

Anatomical images were segmented using Freesurfer 7 [Fischl 2012] to obtain subject-specific anatomical parcellations.

Coregistration of the digitized head position and the reconstructed structural surfaces was performed separately for each session. First the three fiducial points were matched, followed by an iterative closest-point match algorithm that minimizes the distance between the digitized head shape and the skin surface. For the two participants without MRI this procedure consisted in warping the Freesurfer template to match their digitized head shape and derive a subject-specific template. As recently demonstrated, using a template produces highly similar results than using the subject-specific T1 image [Vinding et al., 2022].

A single shell boundary-element method model was created to define a volume source space by filling the inner skull surface with equidistant (5 mm) voxels.

Inverse solution was based on linearly-constrained miminum variant (LCMV) beamformer [Van Veen et al., 1997]. A beamformer approach was chosen given its better resolution in estimating subcortical activity [Ruzich 2019; Pu 2018]. A time domain solution was preferred to be able to reconstruct multiple frequencies at source level using a single spatial filter, such that the observed differences cannot be ascribed to differences in the inversion algorithm. Artifact-free segments were used to estimate the data covariance (separately for the 15° and 30° conditions) while subject specific empty-room recordings were used to model noise and account for the different contributions of the two sensor types. Both data and noise covariance matrices were regularized with 5% of the sensors’ power and their rank was reduced to the residual degrees of freedom after application of MaxFilter [Westner 2022]. We further reduced the dimensionality of the inversion kernel by retaining (through SVD) the dipole orientation that maximized power resulting in a scalar beamformer. The FFT and then ITC were computed at each voxel from the reconstructed time series as detailed in the frequency analysis section.

## QUANTIFICATION AND STATISTICAL ANALYSIS

### Behavioural analysis

Accuracy (percentage of correct responses) was computed separately for each session and each angular resolution. We used a dependent samples t-test to compare these metrics across resolutions, separately for each session. One participant of the spatial experiment was excluded from further analysis due to poor performance (mean accuracy: dots: 54%; lines: 62%).

### Sensor-level cluster-permutation test

A two-sided cluster-based permutation test [Maris & Oostenveld 2007] was used to compare the ITC at the frequency corresponding to the 60° spatial periodicity (ITC_60°,_1.5 Hz in the 15° resolution, 3 Hz in the 30° resolution) to the ITC of the control spatial periodicities (15° resolution: ITC_45°_ 2 Hz, ITC_90°_ 1 Hz; 30° resolution: ITC_90°_ 2 Hz), separately for each angular resolution and each control periodicity. For this analysis we considered only magnetometers, given their higher sensitivity to deep brain structures as compared to planar gradiometers [Hari 2012].

In brief, a one-sample t-test is performed at each channel on the difference between conditions (i.e., spatial periodicities). The channels that survived an uncorrected threshold of p < 0.05 are retained to form spatial clusters based on a predefined adjacency matrix with ~ 6 neighbours per channel. This procedure was repeated 10000 times, each time shuffling the condition labels and retaining the highest cluster statistic (t-score). A p-value corrected for multiple comparisons is obtained by comparing the cluster statistic observed from the actual contrast with the distribution of permuted cluster statistics.

The same analysis was applied to the non-spatial experiment data, comparing the frequencies that in the spatial experiment were tagged with spatial regularities.

### Source-level ROI analysis

Source level analysis focused on subject-specific anatomical regions of interest (ROI) obtained from the Freesurfer parcellation. Specifically, we created a medial-temporal lobe (MTL) ROI encompassing the entorhinal and parahippocampal cortex, from the Desikan-Killiany atlas [Desikan et al., 2006], and the hippocampus, obtained from Freesurfer’s own subcortical parcellation [Fischl et al., 2002]. As control ROIs we used the lateral occipital and precentral regions from the Desikan-Killiany atlas.

Subject-specific, average ITC values within each ROI were entered into series of analysis of variance (ANOVA) in R (R Core Team, 2022). First, a two-way repeated measures ANOVA with factors session (dots, lines) x periodicity (15° resolution: 90°, 60°, 45°; 30° resolution: 90°, 60°) was aimed at investigating differences in the neural tracking in the MTL ROI between the dots and lines sessions. This analysis was repeated for each hemisphere and each angular resolution. Results showed no statistically significant session x periodicity interaction, suggesting that a similar response profile could be observed across sessions. All subsequent analysis will therefore consider the average across session as input.

A two-way repeated measures ANOVA with factors ROI (MTL, lateral occipital, precentral) x periodicity (15° resolution: 90°, 60°, 45°; 30° resolution: 90°, 60°) investigated differences across ROIs in the neural tracking of the spatial periodicities. Planned paired t-tests were ran to evaluate whether ITC_60°_ was greater than ITC of the control periodicities in each ROI.

Last, we compared the spatial experiment to the non-spatial experiment. We ran a three-way mixed ANOVA with experiment (spatial, non-spatial) as between-subjects factor and ROI (MTL, lateral occipital, precentral) and periodicity (15° resolution: 90°, 60°, 45°; 30° resolution: 90°, 60°) as within-subject factors. Individual ROIs were further investigated with two-way mixed ANOVAs with factors experiment x periodicity. Greenhouse-Geisser correction was applied to the degrees of freedom in case sphericity assumption was violated.

### Conjunction analysis

ROI analysis is biased by a-priori selection of a limited number of regions, effectively neglecting contributions from other parts of the brain. To assess whether this constraint hinders the interpretation of our findings we additionally computed a whole-brain conjunction analysis [Nichols 2005]. This consists in computing the individual contrasts between ITC_60°_ and ITC of the control periodicities for each cortical voxel and for each angular resolution. The resulting t-maps are then combined by retaining, for each voxel, the minimum t-value across all contrasts and angular resolutions. Resulting t-maps are plotted at an uncorrected threshold of p < 0.005.

### Bayesian model comparison

To directly compare the ITC_60°_ with both control ITCs at the same time, we fitted linear and quadratic models to the group-level ROI data at 15° resolution. For both linear and quadratic models we computed the Bayesian information criterion (BIC), a metric that quantifies goodness of fit while accounting for the number of parameters included in the model. Results were compared using a Bayes Factor (BF) [Wagenmakers 2007]. In brief, BF is computed as follows:

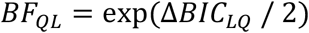

where Δ*BIC*_LQ_ is the difference in the BIC obtained for each model. QL indicates the evidence in favour of the quadratic (Q) over the linear (L) model and vice versa. This formulation is consistent with the “unit information prior”. The BF quantifies the strength of the evidence for one model as compared to the other and can be interpreted according to standard guidelines [Raftery 1995].

### Topographies’ correlation between experiments and spatial periodicities

Having observed a similar, significant frequency preference in both experiments in the 30° resolution, we sought to quantify the extent to which this similar response is expressed in the pattern of sensor level activity. To this end, we correlated the group-average topography of the ITC difference between frequencies tagged with 60° and 90° periodicity between the spatial (i.e., Fig. 2C, right) and non-spatial experiment (i.e., Fig. 4A, right) in the 30° resolution.

Moreover, we reasoned that a grid-like response should be relatively independent of the granularity of the space that is used in its investigation, thus providing a similar response pattern in both spatial resolutions that were tested in the current experiment. To investigate this we computed the (within-participant) similarity of the topographies of the grid-like effect (i.e., ITC_60°_ - ITC_90°_) between the 15° and 30° angular resolutions. We focused on 60° and 90° periodicities given that they were common between angular resolutions. Similarly, in the non-spatial experiment we computed the correlation of the topographies (ITC_Cont60°_ - ITC_Cont90°_) between the conditions corresponding to the 15° and 30° resolutions. Correlation values were fisher transformed before further analysis. A one-sample t-test was used to investigate whether the correlations at the group level were significantly different from zero. An independent samples t-test was instead used to compare the correlation scores across groups.

