## Supplementary results for "MEG frequency tagging reveals a grid-like code during covert attentional movements"

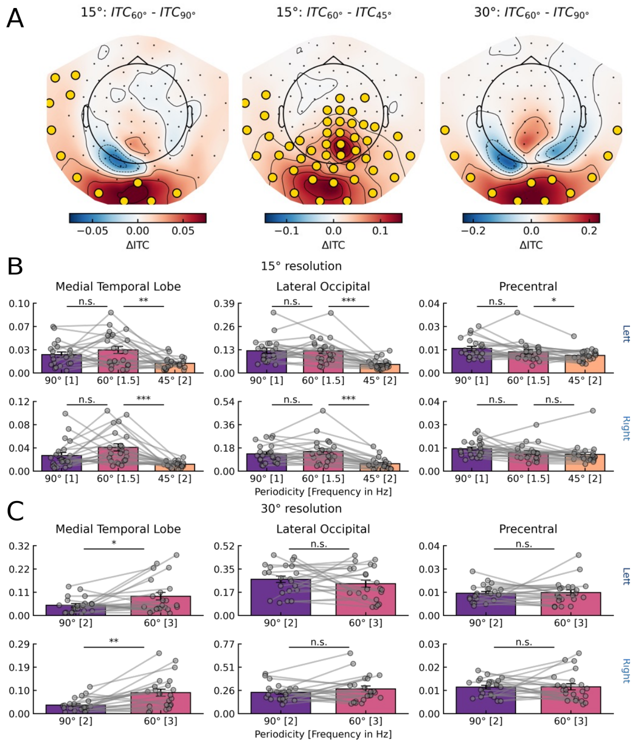


Figure S1 Sensor- and source-level results for the dots session.

A) Significant clusters at sensor-level in which ITC60° is greater than control ITCs. In the 15° resolution ITC60° is greater than ITC90° (left, p=0.009 cluster corrected) and ITC60° is greater than ITC45° (center, p<0.001). In the 30° resolution ITC60° is greater than ITC90° (right, p=0.011). No clusters were found in which control periodicities are higher than 60° periodicity.

B) A two-way repeated measures ANOVA with factors periodicity (90°, 60°, 45°) and ROI (MTL, lateral occipital, precentral) identified significant interactions in both the left (top, F(4, 84)=13.49, p<0.001) and right hemisphere (bottom, F(4, 84)=13.667, p<0.001). Planned paired t-test did not identify significant differences between ITC60° and ITC90° in neither the left (t(21)=1.14, p=0.266) nor the right hemisphere (t(21)=1.70, p=0.102) but a significantly greater ITC60° than ITC45° in both the left (t(21)=3.31, p=0.003) and the right hemisphere (t(21)=4.49, p<0.001). In the lateral occipital no significant differences were identified between ITC60° and ITC90° in neither the left (t(21)=-0.07, p=0.944) nor the right hemisphere (t(21)=1.01, p=0.323) but a significantly greater ITC60° than ITC45° in both the left (t(21)=4.02, p<0.001) and the right hemisphere (t(21)=6.35, p<0.001). In the precentral ROI we did not find significant differences between ITC60° and ITC90° in neither the left (t(21)=-1.25, p=0.223) nor the right hemisphere (t(21)=-1.48, p=0.151) but a significantly greater ITC60° than ITC45° in the left (t(21)=2.09, p=0.048) but not the right hemisphere (t(21)=1.23, p=0.230).

C) A two-way repeated measures ANOVA with factors periodicity (90°, 60°) and ROI (MTL, lateral occipital, precentral) identified significant interactions in the left (top, F(2, 42)=3.40, p=0.042) but not in the right hemisphere (top, F(2, 42)=2.20, p=0.123). However, planned paired t-tests identify a significantly greater ITC60° as compared to ITC90° in both the left (t(21)=2.25, p=0.034) and right hemisphere (t(21)=3.62, p=0.001). No significant differences between the ITCs were found in any control region (left: lateral occipital: t(21)=-0.93, p=0.361; precentral: t(21)=0.21, p=0.831; right: lateral occipital: t(21)=1.14, p=0.263; precentral: t(21)=0.09, p=0.922).

Gray dots indicate individual subjects. Error bars indicate standard error of the mean. Lines above data points indicate significance (n.s.: not significant; *: p < 0.05; **: p < 0.01; ***: p < 0.001)

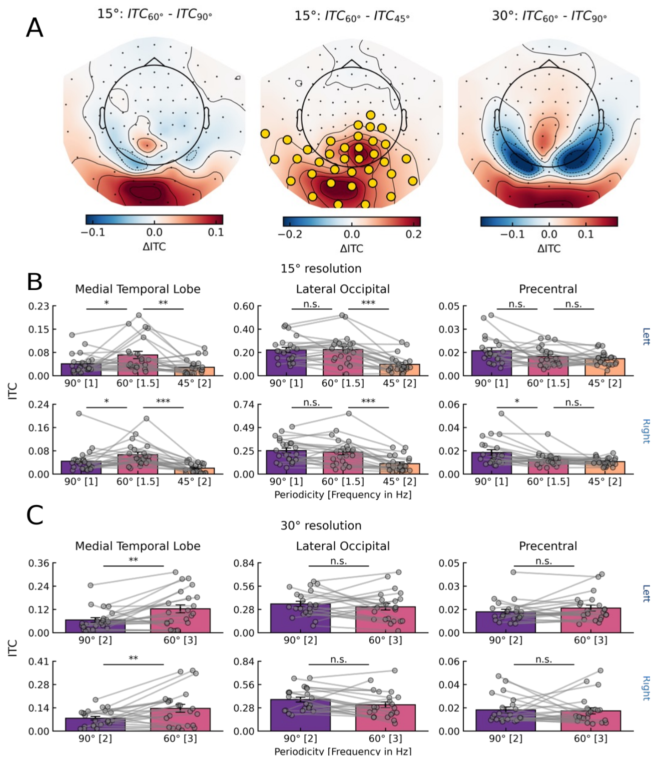


Figure S2 Sensor- and source-level results for the lines session.

A) Cluster-permutation test at sensor-level identifies a significant cluster in which ITC60° is greater than ITC45° (p=0.007 cluster corrected). No significant clusters were found in the other contrasts (all p>0.1), although topographies were very similar to the dots session.

B) A two-way repeated measures ANOVA with factors periodicity (90°, 60°, 45°) and ROI (MTL, lateral occipital, precentral) identified significant interactions in both the left (top, F(4, 84)=26.15, p<0.001) and right hemisphere (bottom, F(4, 84)=17.27, p<0.001). Planned paired t-test identified significant differences between ITC60° and ITC90° in the left (t(21)=2.39, p=0.026) and the right hemisphere (t(21)=2.17, p=0.040) as well as a significantly greater ITC60° than ITC45° in both the left (t(21)=3.13, p=0.005) and the right hemisphere (t(21)=5.39, p<0.001). In the lateral occipital no significant differences were identified between ITC60° and ITC90° in neither the left (t(21)=0.19, p=0.848) nor the right hemisphere (t(21)=-0.78, p=0.439) but a significantly greater ITC60° than ITC45° in both the left (t(21)=5.79, p<0.001) and the right hemisphere (t(21)=5.26, p<0.001). In the precentral ROI we did not find significant differences between ITC60° and ITC90° in the left (t(21)=-1.77, p=0.089) but a significant difference in the right hemisphere (t(21)=-2.62, p=0.015) while no significant differences were found between ITC60° than ITC45° neither in the left (t(21)=1.00, p=0.325) nor in the right hemisphere (t(21)=1.14, p=0.267).

C) A two-way repeated measures ANOVA with factors periodicity (90°, 60°) and ROI (MTL, lateral occipital, precentral) identified significant interactions in the left (top, F(2, 42)=4.15, p=0.023) and in the right hemisphere (top, F(2, 42)=10.78, p<0.001). Planned paired t-tests identify a significantly greater ITC60° as compared to ITC90° in both the left (t(21)=3.21, p=0.004) and right hemisphere (t(21)=2.84, p=0.009). No significant differences between the ITCs were found in any control region (left: lateral occipital: t(21)=-0.83, p=0.413; precentral: t(21)=1.44, p=0.162; right: lateral occipital: t(21)=-1.92, p=0.067; precentral: t(21)=0.23, p=0.817).

Gray dots indicate individual subjects. Error bars indicate standard error of the mean. Lines above data points indicate significance (n.s.: not significant; *: p < 0.05; **: p < 0.01; ***: p < 0.001)

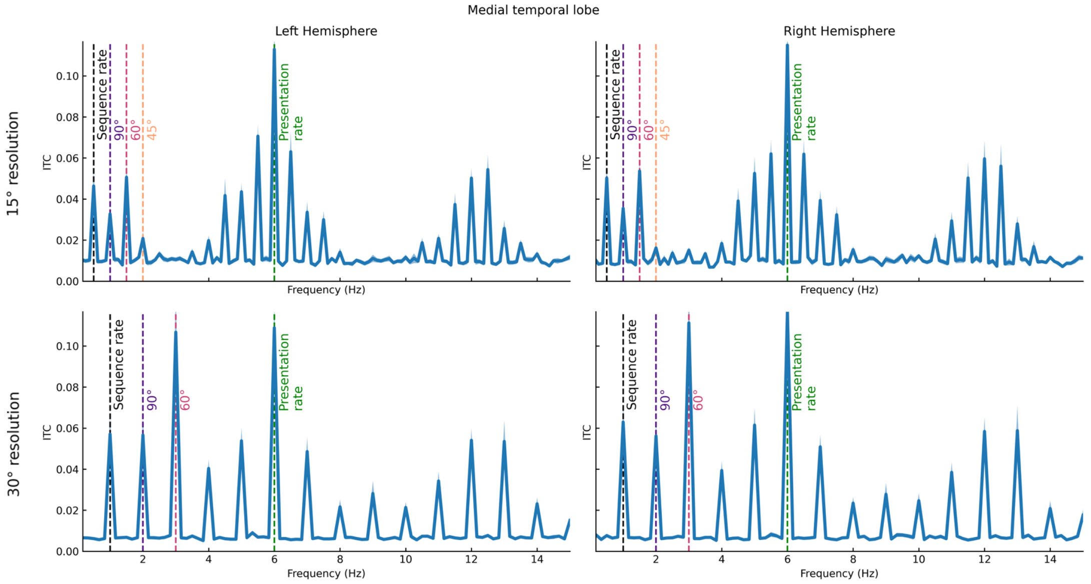


Figure S3 Intermodulation effect in the spatial experiment.

Multiple peaks were present in the frequency spectrum (0.1-15 Hz) in the MTL ROI (left column: left hemisphere; right column: right hemisphere), not only the presentation rate and the ones tagged with spatial regularities. The lowest (in Hz) prominent peak was the one at the "sequence rate", i.e., the time interval at which an individual sequence of unique trajectories was repeatedly presented (0.5 Hz in 15° resolution (top), and 1 Hz in 30° resolution). In other words, the time interval to complete one "turn" of the clock-like presentation. Note that one sequence (or turn) corresponded to half of the clock, given the lack of directionality that made opposite trajectories indistinguishable: e.g. a trajectory starting at 20° and ending at 200° occupied the same portion of space than a trajectory starting at 200° and ending at 20°. The peaks corresponding to the spatial regularities (highlighted) could then be reinterpreted as being the harmonics of this "sequence rate" peak or the intermodulation between the "sequence rate" and the presentation rate. Intermodulations are in fact any sum of the multiples of the original frequencies, e.g. considering 30° resolution 1 Hz (f1, sequence rate) and 6 Hz (f2, presentation rate), intermodulation components can arise at f1+f2=7Hz, f2-f1=5Hz, f2-f1*2=4Hz, f2*2-f1=11Hz etc.. Although the frequency tagged with 60° periodicity was the one with the highest ITC (excluding presentation rate), we could not rule out the possibility that its ITC was a byproduct of these intermodulation components. In fact, intermodulation components can give rise to a frequency spectrum similar to the one we found in the MTL (see Gordon et al., 2019 Fig. 2). This finding motivated us to run a control experiment to investigate the effect that temporal regularities corresponding to a sequence rate have on the neural response.

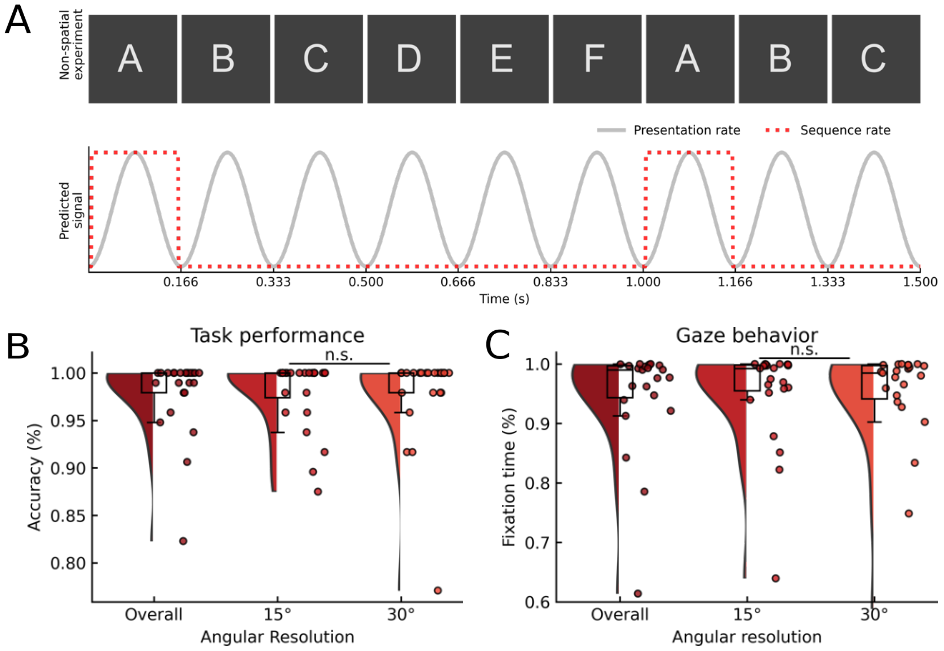


Figure S4 Design of the non-spatial experiment and behavioral results.

A) Example of a sequence of letters corresponding to the 30° resolution. Individual letters were presented continuously every 167 ms (6 Hz). This same sequence of letters (A to F) was presented repeatedly every 1 s (1 Hz), generating a rhythm corresponding to the temporal structure of the sequence that corresponds to the presentation of a 30° sequence in the spatial experiment. Similarly, the 15° condition is mimicked by creating a sequence of 12 letters (A to N) that was repeated every 2 s (0.5 Hz) corresponding to the time required to present a sequence of trajectories linearly-spaced by 15° in the spatial experiment. With this presentation no frequency is tagged with spatial periodicity. The frequencies of interest of the spatial experiment (e.g., 2 and 3 Hz in 30°) in fact correspond to the presentation of letters A and D and A and E, respectively.

B) Accuracy in the color-change detection task, averaged over angular resolutions as well as separately for each angular resolution. Participants were overall accurate (M=97.87%, SD=4.02%) except for one participant that was 2SD below the group mean and was thus excluded from further analyses. No difference was found between conditions ($t(23)=0.13$, $p=0.896$).

C) Fixation time during the trials, expressed as percentage, averaged over angular resolutions as well as separately for each angular resolution. Participants were keeping fixation within a 4.5° fixation window for the majority of the time during the trial. Trials in which fixation was maintained for less than 80% of the time were excluded from further analyses and this led to the exclusion of one participant due to the low amount of trials left.

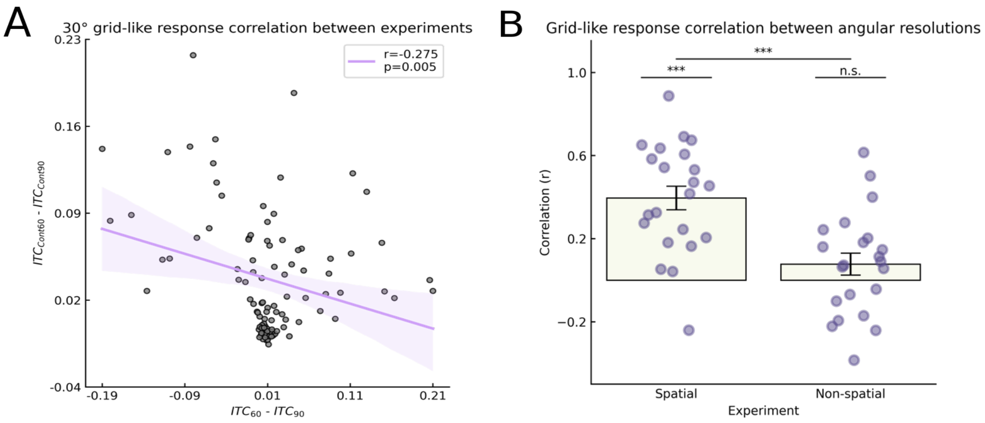


Figure S5 Topographies correlation between experiments.

A) Although the cluster-permutation test identified similar frequency preferences in the spatial and non-spatial experiment, in the 30° condition (see Fig 2C and 4A in main text), topographies seemed quite different, suggesting that these frequency preferences may rely on different neural mechanisms. To directly assess the similarity of the neural response between the experiments, we correlated the group-level topography of the grid-like response in the 30° condition between the spatial and non-spatial experiment and found a significant negative correlation ($r(100)=-0.227$, $p=0.005$) suggesting differences in the neural mechanism generating the measured response, even if a significant difference in the frequencies of interest is found in both experiments.

B) We reasoned that the spatial resolution that defines the space should not influence the grid-like response, thus if grid cells are contributing to the signal we are measuring we should observe a high correlation between the grid-like response in the 15° and 30° resolution, but only in the spatial experiment. To test this hypothesis, we measured the (within-subject) similarity between the multivariate grid-like response (i.e. the difference between the tagged frequencies that are in common between the two resolutions) in the 15° and 30° resolution in both the spatial and non-spatial experiment. Purple dots indicate individual subjects. Error bars indicate standard error of the mean. Lines above data points indicate significance (n.s.: not significant; *: $p < 0.05$; **: $p < 0.01$; ***: $p < 0.001$) Confirming the prediction, we found a significant correlation at the group level between the 15° and 30° topographies in the spatial experiment ($t(21)=6.09$, $p<0.001$) and not in the non-spatial experiment ($t(21)=1.48$, $p=0.151$), with the correlation in the spatial experiment being higher than in the non-spatial experiment (independent samples $t(42)=3.99$, $p<0.001$).

These results indicate that although univariate analyses identify comparable differences in the ITC response in both the spatial and non-spatial experiment, the multivariate topographical pattern can provide additional information and distinguish the neural tracking of the spatial periodicities as compared to a response to the temporal regularities.

| Angular Resolution | Hemisphere | ROI | Comparison | df | t | p | Sig. |
|---|---|---|---|---|---|---|---|
| 15° | Left | Mtl | Cont60°[1.5 Hz] vs Cont90°[1 Hz] | 21 | 1.74 | 0.094 | n.s. |
| | | | Cont60°[1.5 Hz] vs Cont45°[2 Hz] | 21 | -1.41 | 0.171 | n.s. |
| | | Lat. occipital | Cont60°[1.5 Hz] vs Cont90°[1 Hz] | 21 | 1.65 | 0.112 | n.s. |
| | | | Cont60°[1.5 Hz] vs Cont45°[2 Hz] | 21 | -6.09 | <0.001 | *** |
| | | Precentral | Cont60°[1.5 Hz] vs Cont90°[1 Hz] | 21 | 2.04 | 0.053 | n.s. |
| | | | Cont60°[1.5 Hz] vs Cont45°[2 Hz] | 21 | -2.85 | 0.009 | ** |
| | Right | Mtl | Cont60°[1.5 Hz] vs Cont90°[1 Hz] | 21 | 0.73 | 0.468 | n.s. |
| | | | Cont60°[1.5 Hz] vs Cont45°[2 Hz] | 21 | -2.70 | 0.013 | * |
| | | Lat. occipital | Cont60°[1.5 Hz] vs Cont90°[1 Hz] | 21 | 4.83 | <0.001 | *** |
| | | | Cont60°[1.5 Hz] vs Cont45°[2 Hz] | 21 | -1.61 | 0.121 | n.s. |
| | | Precentral | Cont60°[1.5 Hz] vs Cont90°[1 Hz] | 21 | -0.45 | 0.652 | n.s. |
| | | | Cont60°[1.5 Hz] vs Cont45°[2 Hz] | 21 | -3.19 | 0.004 | ** |
| 30° | Left | Mtl | Cont60°[3 Hz] vs Cont90°[2 Hz] | 21 | 3.40 | 0.002 | ** |
| | | Lat. occipital | Cont60°[3 Hz] vs Cont90°[2 Hz] | 21 | 3.49 | 0.002 | ** |
| | | Precentral | Cont60°[3 Hz] vs Cont90°[2 Hz] | 21 | 0.76 | 0.450 | n.s. |
| | Right | Mtl | Cont60°[3 Hz] vs Cont90°[2 Hz] | 21 | 2.28 | 0.032 | * |
| | | Lat. occipital | Cont60°[3 Hz] vs Cont90°[2 Hz] | 21 | 1.13 | 0.267 | n.s. |
| | | Precentral | Cont60°[3 Hz] vs Cont90°[2 Hz] | 21 | 0.88 | 0.388 | n.s. |

Table S1. Pairwise comparison between frequencies at source level in the non-spatial experiment.
We ran post-hoc t-tests at source level in each ROI to investigate whether a frequency preference was present in the non-spatial experiment (see Fig. 3 for the same tests in the spatial experiment).
In the condition corresponding to 15° resolution (top), in the left hemisphere we found no significant difference between the ITCs ($ITC_{Cont60°}$ vs $ITC_{Cont90°}$: t(21)=1.74, p=0.094; $ITC_{Cont60°}$ vs $ITC_{Cont45°}$: t(21)=-1.41, p=0.171) while in the right we found a significant difference between $ITC_{Cont60°}$ and $ITC_{Cont45°}$ (t(21)=-2.70, p=0.013) and no significant difference between $ITC_{Cont60°}$ and $ITC_{Cont90°}$ (t(21)=0.73, p=0.468). In control regions Post-hoc t-tests in the left lateral occipital show that $ITC_{Cont45°}$ was greater than $ITC_{Cont60°}$ (t(21)=-6.09, p<0.001) while no significant difference was found between $ITC_{Cont60°}$ and $ITC_{Cont90°}$ (t(21)=1.65, p=0.112). The opposite effect was found in the right lateral occipital, with $ITC_{Cont60°}$ being greater than $ITC_{Cont90°}$ (t(21)=4.83, p<0.001) but no significant difference between $ITC_{Cont60°}$ and $ITC_{Cont45°}$ (t(21)=-1.61, p=0.121). In the left precentral ROI instead we found that $ITC_{Cont45°}$ was significantly higher than $ITC_{Cont60°}$ (t(21)=-2.85, p=0.009) while no significant difference was found between $ITC_{Cont60°}$ and $ITC_{Cont90°}$ (t(21)=2.04, p=0.053). Similarly in the right hemisphere only $ITC_{Cont45°}$ was greater than $ITC_{Cont60°}$ (t(21)=-3.19, p=0.004) but no significant difference was found between $ITC_{Cont60°}$ and $ITC_{Cont90°}$ (t(21)=-0.45, p=0.652).
In the condition corresponding to the 30° resolution (bottom), post-hoc t-tests in the MTL revealed a significantly greater $ITC_{Cont60°}$ than $ITC_{Cont90°}$ (left: t(21)=3.40, p=0.002; right: t(21)=2.28, p=0.032). In the right hemisphere we found no significant difference between the ITCs in neither control ROI (lateral occipital: t(21)=1.13, p=0.267, precentral: t(21)=0.88, p=0.388). Post-hoc t-tests in the left hemisphere demonstrate that, in the lateral occipital, $ITC_{Cont60°}$ was greater than $ITC_{Cont90°}$ (t(21)=3.49, p=0.002) while no difference was found in the precentral ROI (t(21)=0.76, p=0.450).
